# DNA damage-induced lncRNA MEG9 impacts angiogenesis

**DOI:** 10.1101/2022.12.07.519382

**Authors:** Eugenia Fraile-Bethencourt, Sokchea Khou, RaeAnna Wilson, Adrian Baris, Rebecca Ruhl, Cristina Espinosa-Diez, Sudarshan Anand

## Abstract

Endothelial cells are highly responsive to environmental changes that allow them to adapt to intrinsic and extrinsic stimuli and switch their transcriptome accordingly to go back to vascular homeostasis. Our previous data demonstrated that small non-coding-RNAs respond quickly to genotoxic stressors and determined endothelial cell fate and DNA damage response. To further understand the contribution of non-coding-RNAs, we profiled differentially expressed long non-coding RNAs in response to genotoxic stress and compared them to pro-angiogenic growth factor signaling. We identified the Maternally expressed gene 9 (MEG9) as a cytoprotective lncRNA in the endothelium. Gain and Loss-of-function studies indicate that MEG9 prevents endothelial cells from cell death, suggesting that MEG9 responses to genotoxic stress can be an adaptive and protective mechanism. Consistent with this phenotype, the knockdown of MEG9 decreases growth factor-dependent angiogenesis in a 3D fibrin gel angiogenesis assay. Deletion of the MEG9 ortholog, Mirg, in mice results in increased vascular leak in Matrigel plugs and a sex and age-dependent decrease in platelets. Mechanistically, we observed that both MEG9 knockdown in vitro and *Mirg*-deleted mice in vivo activated common pathways, including apoptosis, clotting, and inflammation. Indeed, the proinflammatory adhesion molecule ICAM1 was significantly increased in human and mouse endothelial cells in a MEG9-dependent manner, supporting the increased vascular permeability observed on MEG9 deficient cells. Taken together, our findings illustrate how genotoxic stress responses through dynamic modulation of lncRNAs, such as MEG9, trigger adaptive mechanisms to maintain endothelial function, while loss of these molecules contributes to maladaptive responses and endothelial cell dysfunction.

## Introduction

Blood vessel cells are susceptible to environmental changes, including mechanical (blood flow, pressure, shear stress) or biochemical (oxygen, oxidative stress, genotoxic, inflammation). The endothelial cell monolayer is often the most exposed and the fastest to respond accordingly to specific stressors or stimuli. Many of these responses are also exquisitely context- and tissue-dependent across different vascular niches. We and others have described the role of non-coding RNAs in distinct endothelial stress responses across pathologies [1-7]. We previously described a signature of genotoxic-stress-induced microRNAs that can control the formation of new vessels, angiogenesis, in the context of tumor microenvironment by influencing endothelial cell fate decisions from senescence to apoptosis. [5, 7].

However, the role of long-non-coding-RNAs in response to stress in the endothelium and the biological mechanisms by which specific lncRNAs regulate vascular stress responses is not fully appreciated. LncRNAs are non-coding RNAs longer than 200 nucleotides that control gene expression through epigenetic, transcriptional, and post-transcriptional mechanisms but also interact with functional biomolecules due to their secondary structure [8, 9]. LncRNAs are critical regulators of the cardiovascular system from development [10, 11], lineage identity[12], endothelial cell dysfunction to angiogenesis in physiological and pathological conditions [13-18]. Emerging evidence has defined some specific lncRNAs as regulators of DNA damage and senescence in the vasculature [19]. Additionally, the potential utility of lncRNAs as targets for the development of vascular disease therapies is currently of great interest [20, 21].

Several stressors, such as oxidative stress, cytotoxic agents, and radiotherapy are known to cause DNA damage and trigger repair pathways in the vasculature [22]. These stressors cause various molecular changes, including telomere shortening, base damage, adduct formation, mismatch, and double-strand breaks. The failure of the vasculature to repair such damage can lead to mutations, cell cycle arrest, cellular senescence, inflammation, EC dysfunction, and apoptosis. The pathological consequences of these DNA-damaging mechanisms are often associated with vascular disease, especially atherosclerosis. Some studies have identified specific lncRNAs as regulators of DNA damage responses [15, 23-25].

Similarly, while the role of angiogenic growth factors has been well-studied, their interaction with DNA repair pathways still needs to be fully understood [26, 27]. Given our prior work identifying distinct roles for microRNAs during DNA damage and angiogenic growth factor signaling, we postulated that lncRNAs at the intersection of DNA damage and angiogenesis play critical roles in endothelial pathophysiology. Consequently, we undertook a selective lncRNA screening approach to identify the differentially expressed lncRNAs in response to pro-angiogenic growth factors and several defined genotoxic stressors.

We found that the LncRNA MEG9 is significantly upregulated after acute exposure to different genotoxic stressors in endothelial cells, while it is decreased in response to growth factors. This lncRNA is encoded in the non-coding RNA cluster DLK1-DIO3, an imprinted genomic region that encodes not only lncRNAs but microRNAs [28, 29], which is highly conserved across mammals [30]. Both microRNAs and lncRNAs from the DlK1-DIO3 cluster have been previously associated with responses to stress and present a role in cardiac and vascular disease [31]. Although the potential regulation of MEG9 in response to genotoxic stressors and ischemic conditions has been studied, the functional role of MEG9 in the vasculature is unexplored. In this manuscript, we describe how disruption of MEG9 expression *in vitro* and *in vivo* induces endothelial cell dysfunction while leading to a more pro-inflammatory phenotype.

## Materials & Methods

### Cell Culture

HUVECs and HMVECs (Lonza) were cultured in EGM-2 media (Lonza) supplemented with 10% Fetal Calf Serum (Hyclone). HUVECs were treated with Cisplatin (5uM), Etoposide (10uM), Hydrogen Peroxide (500uM), Hydroxyurea (5uM) and were purchased from Sigma. 5-Azacytidine was purchased from Sigma. VEGF (50ng/mL), β-FGF (100ng/mL) were purchased from Peprotech, and rDLL4 (10ug/ml), Jagged-1 (10 ug/ml) were purchased from R&D Biosystems. Cells were incubated at 37°C with 5% CO_2_ for 6 hours. VEGF was purchased from PeproTech, Inc. RNA GapmeRs were purchased from Exiqon or Qiagen.

### Transfection

Cells were transfected at 60–70% confluence using standard forward transfection protocols using Lipofectamine RNAimax reagent (Life Technologies) for siRNAs or GapmeRs. Typically, 50 nM RNA was used for transfections following the manufacturer’s protocols.

### RNA extraction and qRT-PCR

Total RNA was isolated using a miRVana microRNA isolation kit (Ambion). Reverse transcription was performed using TaqMan™ Advanced cDNA Synthesis Kit (Life Tech) according to the manufacturer’s instructions. qRT-PCR was performed using multiplexed TaqMan primers (Applied Biosystems) for specific genes MEG9, and Mirg. The relative quantification of gene expression was determined using the 2^-ΔΔCt^ method [32]. Using this method, we obtained the fold changes in gene expression normalized to an internal control gene, GAPDH or U6 snRNA, respectively.

### LncRNA Screen

RNA was diluted to 400ng/uL and cDNA synthesized in duplicate using the Systems Biology LncRNA profiler kit. The resulting cDNA was used in a SYBR-Green based qRT-PCR to analyze the expression of 90 lncRNAs and 6 normalization genes (in duplicate). The data was analyzed, and Ct-values internally normalized to RNU43 to generate ΔCt-values, then treatment conditions were normalized to untreated (IgG treated) cells to generate ΔΔCt-values. Gene expression levels were converted to base-2 fold-change (2^-ΔΔCt^) [32], plotted and compared across conditions.

### Radiation of Cells

Cells were irradiated on a Shepherd 137cesium irradiator at a rate of ∼166 cGy/min.

### Caspase-3/7 Activity & Cell Viability

Cells were transfected with GapMers. After 48 hours, Caspase-3/7 activity and cell viability were measured using Promega Caspase-3/7 Glo and Cell Titer Glo 2.0 according to manufacturer’s protocol.

### BrdU Cell Proliferation Assay

Cells were transfected with GapMers. After 24-hours BrdU was added to the media and 16h later cell proliferation was evaluated using the BrdU Cell Proliferation ELISA kit (EMD-Millipore). Colorimetric analysis was done with single channel 450nm intensity. Blank wells with either media alone or cells without BrdU were used for background correction.

### Western blot and densitometric analysis

After treatment, cells were washed in phosphate-buffered saline (PBS) and lysed in RIPA buffer (Sigma) supplemented with Complete Protease inhibitor cocktail (ROCHE) and Phosphatase inhibitors cocktail 2 and 3 (Sigma). Lysed cells were harvested by scraping, and proteins were analyzed by BCA Assay (Pierce). For protein array experiments, the human apoptosis array (R&D Biosystems ARY009) was used according to the manufacturer’s instructions.

### 3-D Angiogenic Sprouting Assay

Early passage HUVECs were coated on Cytodex-3 beads (GE Healthcare) at a density of 10 million cells/40 μl beads and incubated in suspension for 3-4 hours with gentle mixing every hour. They were plated on TC-treated 6 well dishes overnight and resuspended in a 2mg/ml fibrin gel with 200,000 human vascular smooth muscle cells. The gel was allowed to polymerize, and complete EGM-2 media was added. Sprouts were visualized from days 3-4 via confocal imaging after overnight incubation with FITC labeled Ulex europaeus lectin (Vector labs). Immunofluorescence imaging was performed on a Yokogawa CSU-W1 spinning disk confocal microscope with 20x, 0.45 Plan Fluor objective (Nikon).

#### In vivo methods

All animal work was approved by the OHSU Institutional Animal Use and Care Committee. WT (C57Bl/6N) and Mirg-/-littermates injected subcutaneously with Growth factor-reduced Matrigel BD) with 400 ng mL^-1^ recombinant human bFGF (Millipore). After 7 days Matrigel plugs, as well as heart, lung, liver, spleen, kidneys, and brains, were harvested from mice, and RNA was isolated using the Eurx RNA purification kit according to the manufacturer’s instructions. Matrigel plugs were homogenized and analyzed for hemoglobin content using a colorimetric assay kit (Sigma). In addition, RNA from tissue was used to analyze endothelial activity using the Qiagen endothelial cell activity qRT-PCR array according to the manufacturer’s instructions. Complete blood counts were performed from blood draws from the tail vein or post-mortem cardiac puncture. Blood was collected into EDTA tubes and analyzed using a hematology analyzer (Scivet ABC).

### Statistics

All statistical analysis was performed using Excel (Microsoft) or Prism (GraphPad). Two-tailed Student’s *t*-test or Mann–Whitney *U*-test was used to calculate statistical significance. Data that was not normally distributed as assessed by Shapiro-wilk test (Excel, Real statistics add-in) was evaluated using *U*-test. Variance was similar between treatment groups. A *P* value<0.05 was considered to be significant. Comparison among multiple groups was performed by one-way ANOVA followed by a post hoc test (Tukey’s or Holm-Sidak). In the absence of multiple comparisons, Fisher’s LSD test was used.

## Results

### MEG9 is a stress-induced and growth factor-suppressed lncRNA in endothelial cells

Several publications have previously described the increasing role of lncRNAs in physiological and pathological angiogenesis [33-35]. We previously characterized several miRs involved in endothelial responses to angiogenic growth factors, radiation, genotoxic stress, and ER stress[5, 7]. With the goal of identifying lncRNAs that are regulators of similar stress and growth factor responses, we performed a focused qRT-PCR array screen for lncRNAs annotated in the human genome. We treated Human Umbilical Vein Endothelial Cells (HUVECs) with three distinct stressors – Genotoxic stress induced by etoposide, replication stress induced by hydroxyurea, and oxidative stress induced by hydrogen peroxide (Fig 1A). Similarly, we treated HUVECs with distinct pro-angiogenic stimuli – VEGF, bFGF, and ligands of the Notch pathway DLL4 and Jagged-1 (Fig 1B). By comparing median ranks across the treatment groups, we identified MEG9 as the 6^th^ most upregulated lncRNA by the stressors and the 9^th^ most downregulated lncRNA by the stimuli. Interestingly, MEG9 was among the few lncRNAs that were differentially regulated by stressors vs stimuli. We further validated the array data by qRT-PCR from etoposide and another inducer of genotoxic stress, ionizing radiation, and both increased MEG9 expression in HUVECs (Fig 1C). Similarly, we observed a dose-responsive induction of MEG9 in human microvascular endothelial cells (HMVECs) (Fig 1D).

**Figure 1:**
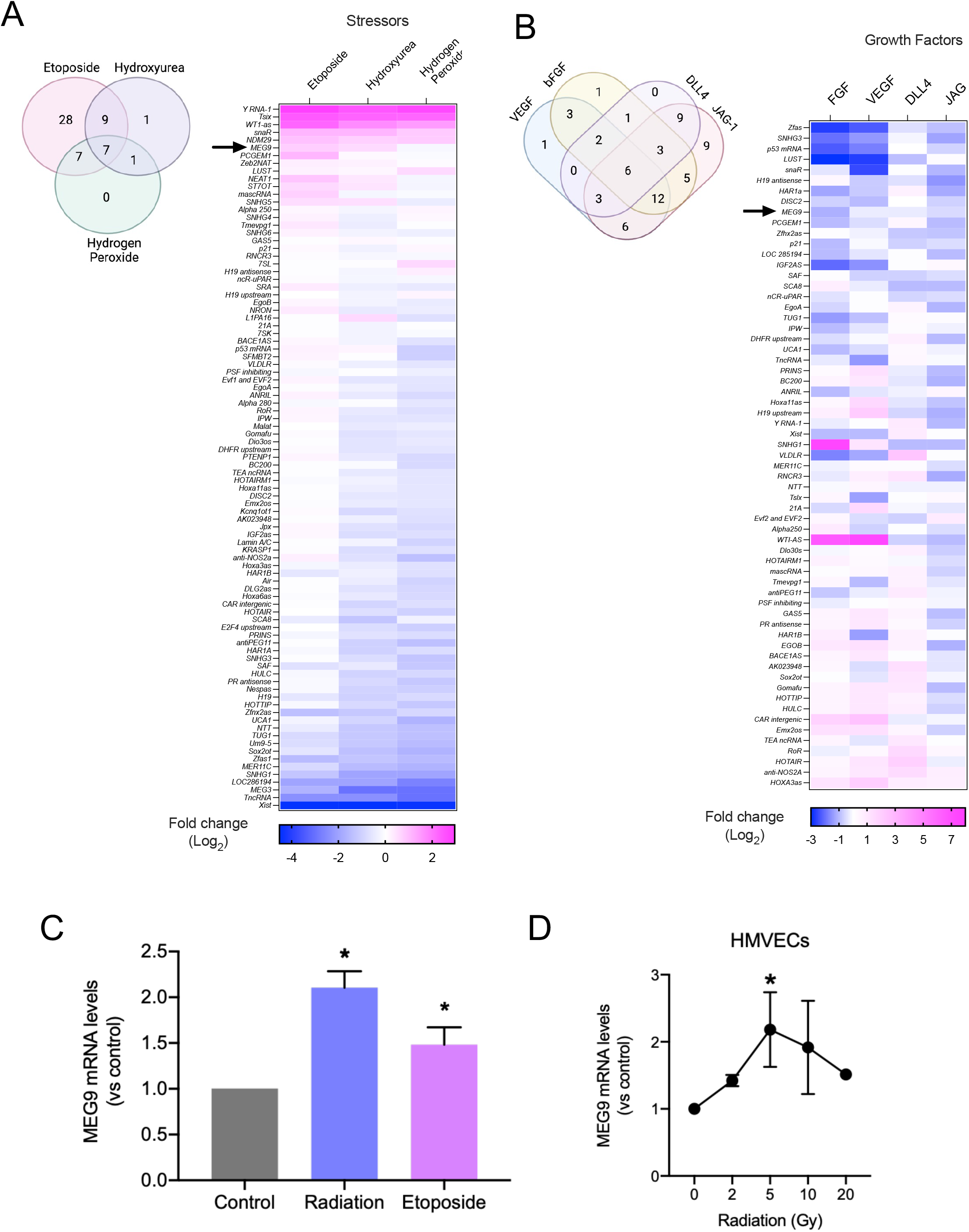
Discovery of MEG9 as a lncRNA induced by DNA damage. HUVECs were starved overnight and treated with indicated stressors -Etoposide (10uM), Hydroxyurea (100uM) or Hydrogen Peroxide (500uM) (A) or growth factors – bFGF (40ug/ml), VEGF (100 ug/ml), DLL4 (10ug/ml) or Jagged-1 (10ug/ml) (B) for 6h. The expression of lncRNAs was measured using a qRT-PCR array for known lncRNAs. C) Validation of MEG9 expression by individual qRT-PCR assay after treatment of HUVECs with radiation (10 Gy) or Etoposide (10 uM). D) Radiation dose-dependent induction of MEG9 in HMVECs. *P<0.05, by Student’s T-test for two groups and ANOVA with post-hoc Sidak test. Heatmaps represent the average of two biological replicates. C-D) Error bars denote S.E.M. of three replicates.

MEG9 is a highly conserved lncRNA across mammals, presenting several isoforms in humans, mouse and cattle [29, 36]. MEG9 is highly enriched in human and mouse brain and other highly vascularized tissues such as lung, liver and heart (Fig 2A-B). Analysis from the Genotype-Tissue Expression (GTEX) database shows MEG9 expression levels across human tissues (Fig 2A) that compares favorably to expression levels in mouse tissues (Fig 2B). Of note, expression levels appeared to diminish with age in the heart and lung tissues (Supplementary Fig 1), something that has been also observed by others in the context of the chronic disease [37]. Interestingly, Single-cell-RNA seq (scRNA seq) data from healthy mouse aorta shows that Mirg expression is fundamentally enriched on the fibroblast populations, although is also expressed in other vascular cells (Supplementary Fig 2A-B)

**Figure 2:**
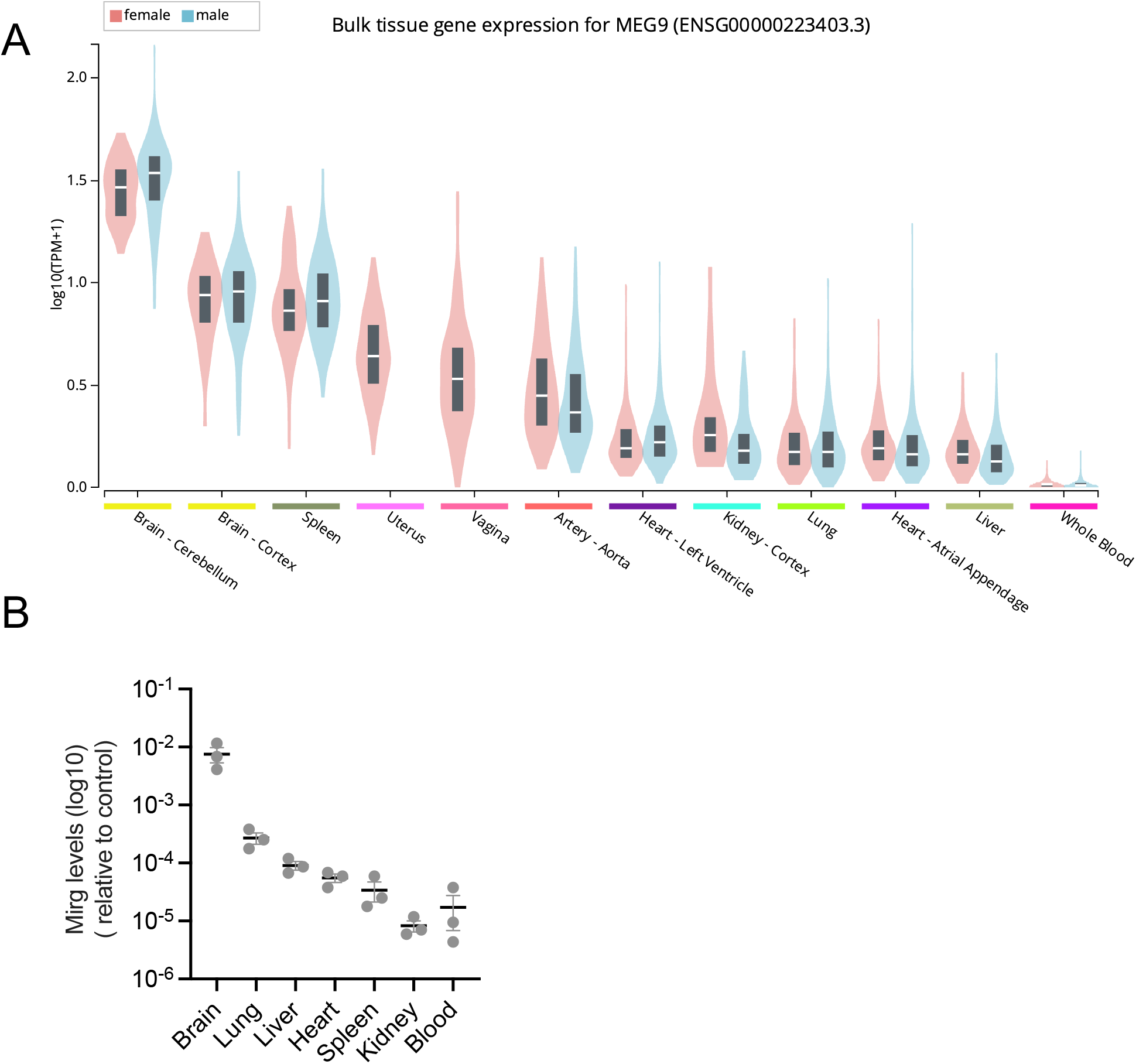
MEG9 expression patterns are homogenous in human and mouse. (A) MEG9 (ENSG00000223403.3) expression in human tissue was obtained from the Genotype-Tissue Expression (GTEx) Project dbGaP Accession phs000424.v8.p2. MEG9 expression is shown in females (pink) and males (blue). (B) Mirg expression was measured by qRT-PCR. RNA was extracted from brain, lung, liver, spleen, kidney, heart and blood.

### Meg9 loss of function leads to endothelial dysfunction in vitro

As MEG9 function in adult endothelial cells has not been described in-depth, we performed a series of gain and loss-of-function experiments. First, we validated that an antisense GapmeR inhibitor significantly decreased MEG9 levels, and that an in vitro transcribed MEG9 RNA approach increased MEG9 levels in HUVECs (Supplementary Fig 3A-B). We found that the MEG9 gain of function decreased active caspase 3 & 7 in decreased serum culture conditions but did not impact cell viability as measured by ATP levels (Fig 3A-B). Indeed, inhibition of MEG9 produced the most robust decrease in cell proliferation as measured by BrdU incorporation compared to a few other lncRNAs with known functions in endothelial cells (Supplementary Fig 4). Consistent with this decrease in proliferation, inhibition of MEG9 also activated apoptotic caspase 3 & 7 pathways and decreased overall viability (Fig 3 C-D). Moreover, inhibition of MEG9 also increased vascular permeability as measured by a FITC-Dextran leak assay (Fig 3E). Finally, to test the impact of these functional changes, we assessed the contribution of MEG9 to angiogenesis in vitro. Consistent with the increase in apoptosis, MEG9 loss-of-function diminished endothelial sprouting in a 3-D fibrin bead angiogenesis assay (Fig 3F). These phenotypic observations indicate that MEG9 might be a pro-survival factor in endothelial cells, the disruption of which causes endothelial dysfunction via cell death, enhanced vascular permeability and decreased angiogenesis.

**Figure 3:**
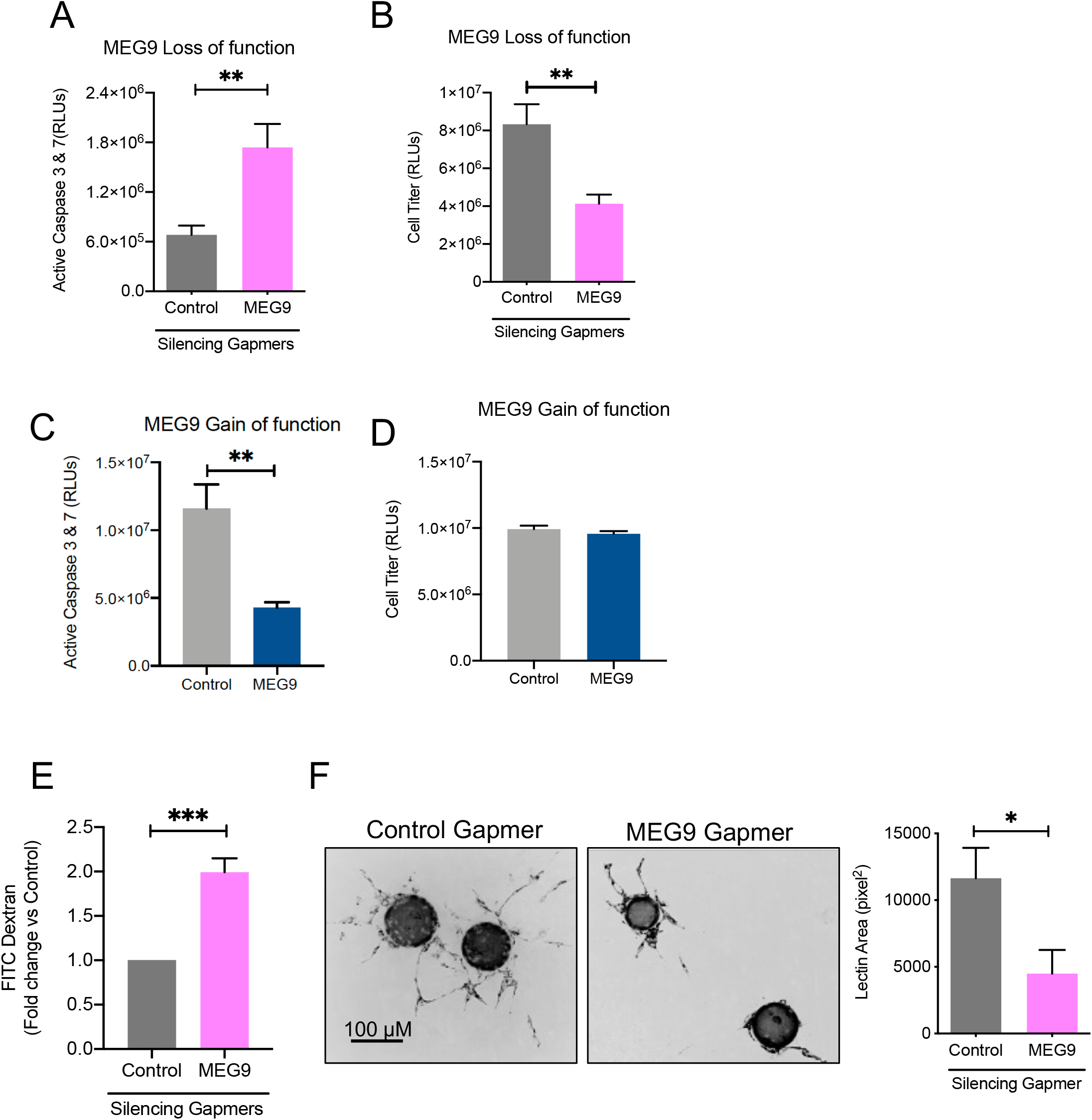
MEG9 inhibition diminishes EC survival and angiogenesis. HUVECs were treated with either control GapmeR or MEG9 GapmeR (A-B) or control RNA or MEG9 RNA (C-D). Cell Death was measured using Caspase 3&7 glo (A, C) and proliferation was measured using Cell Titer glo (B, D) assays. E) Permeability was measured using a FITC-Dextran leak assay. F) Sprouting angiogenesis was measured using a 3D tube formation assay in a fibrin gel. Quantification of sprout area stained using *U*.*europaeus* lectin. Bar graphs show mean + SEM of 20-30 beads per group. One of three independent experiments. *P<0.05, ** P<0.01, *** P<0.005 from a two-tailed Student’s T-test. Bars represent mean ± SEM of independent replicates.

### Loss of the MEG9 ortholog Mirg may impact vascular function in vivo

To evaluate whether the phenotypes we observed in cultured endothelial cells were consistent with the lncRNA function in vivo, we generated a CRISPR-mediated deletion of Mirg in mice targeting a 21kb region encompassing exon1 through exon 8 (Fig 4A, Supplementary Fig 5A-B). Mirg^-/-^ mice were viable, and they did not seem to present any apparent developmental changes or differences in size compared to wild-type littermate controls. First, we confirmed the deletion of Mirg by qRT-PCR (Fig 4A) and observed loss of expression in brain, Matrigel plugs and the hearts of Mirg^-/-^ mice compared to wild-type mice (Fig 4B). Interestingly, while we did not observe any striking phenotype in male Mirg^-/-^ mice, female Mirg^-/-^ mice older than 5 months of age presented an increase in vascular leak in a bFGF Matrigel plug model of angiogenesis (Fig 4C-D) as measured by hemoglobin content. An increase in hemoglobin in this context in a Matrigel plug typically is reflective of increased blood vessels, increased vascular leak, or poor clotting. Therefore, we performed a complete blood count from and found that the female Mirg^-/-^ mice had diminished platelet counts whereas the male mice increased lymphocyte numbers in the peripheral blood (Fig 4E-F). Other cell numbers in the blood were not significantly different from the wild-type mice (Supplementary Figure 6). These observations suggest that lncRNA Mirg may play an important role in mediating thrombosis and vascular leak through endothelial-dependent and independent mechanisms.

**Figure 4:**
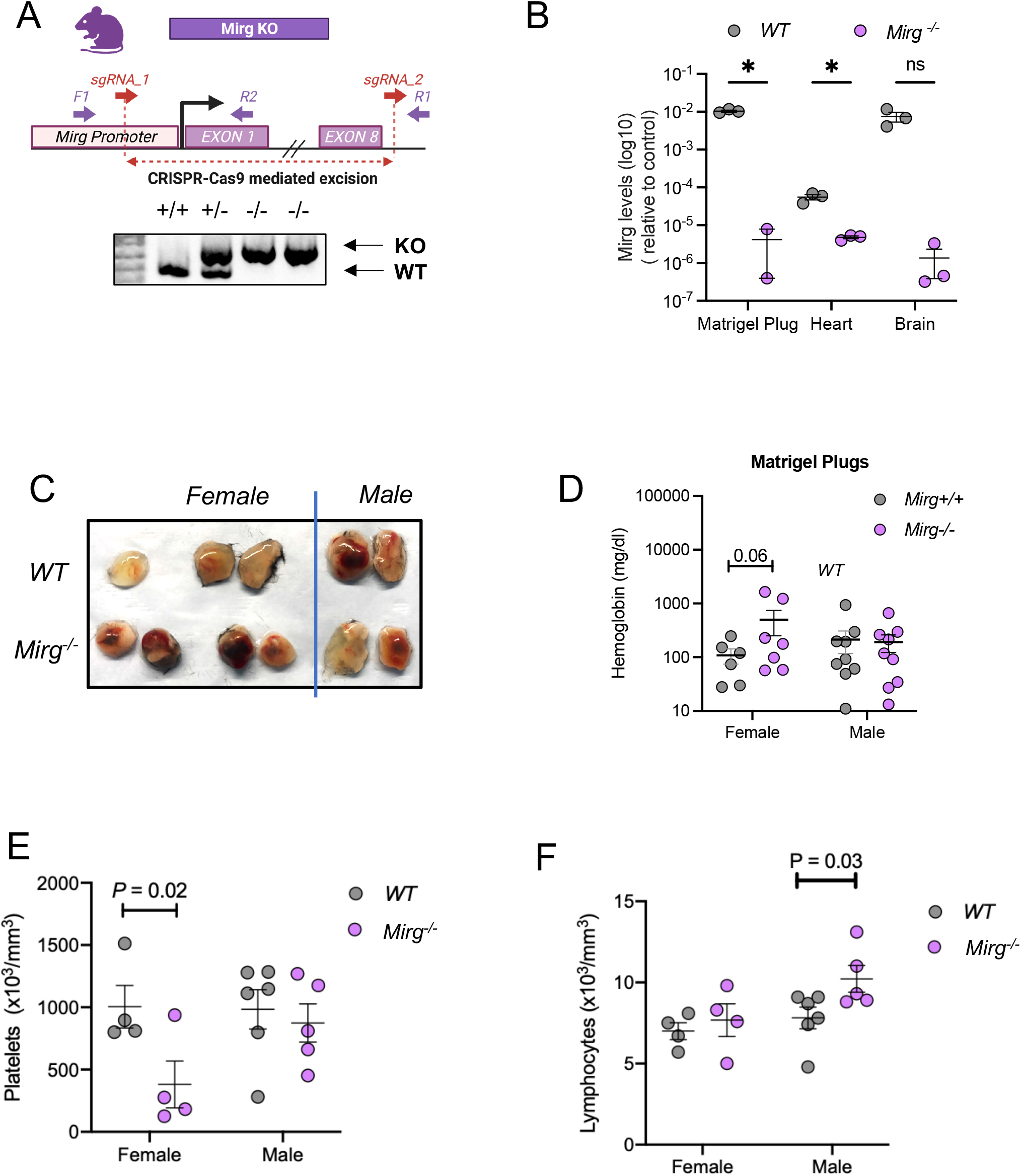
Deletion of mouse MEG9 ortholog Mirg increases vascular leak and decreases platelets in female mice. A) Schematic of Mirg KO CRISPR-CAS9 excision. Genotyping PCR showing efficient deletion on Mirg locus. Mirg^-/-^ or littermate control WT mice were injected with bFGF containing Matrigel on their flank (s.c). One week later, plugs were harvested, and their Mirg levels were quantified using qRT-PCR from the plugs and hearts and brains as representative tissues (B). C-D) Hemoglobin levels were measured from the plugs. E) Platelet and lymphocyte count from Mirg^-/-^ or littermate control WT mice. *P<0.05, ** P<0.01, *** P<0.005 from a two-tailed Student’s T-test.

### Identification of common pathways impacted by human and mouse MEG9

To explore the downstream mechanisms of MEG9 in endothelial cells, we performed a focused qRT-PCR array for endothelial cell function signature from both HUVECs and Mirg^-/-^ mice (Fig 5A-B). We identified several upregulated common pathways, including regulation of apoptosis, angiogenesis, coagulation, and inflammation were significantly impacted by MEG9 or Mirg loss (Fig 5C). Interestingly, integrative analysis of the human and mouse datasets identified two genes-VEGR2 (KDR) and ICAM1 as commonly upregulated in both human and mouse MEG9 loss-of-function models. Similarly, using a western blot-based protein array for angiogenesis pathways, we identified 8 proteins that were dysregulated by MEG9 inhibition (Supplementary Figure 7). Notably, the vasoconstrictor peptide endothelin-1 was significantly downregulated by MEG9 inhibition, while both VEGF and bFGF were modestly downregulated. Among the upregulated proteins, we observed that coagulation factor III was modestly upregulated. However, the involvement of these changes in MEG9’s contribution to endothelial dysfunction remains to be elucidated in future studies. In summary, these observations indicate that MEG9 loss in endothelial cells induces cell death and vascular leak in vitro and in vivo. Moreover, our data indicate that MEG9 might have a non-endothelial cell-autonomous function in thrombosis/coagulation.

**Figure 5:**
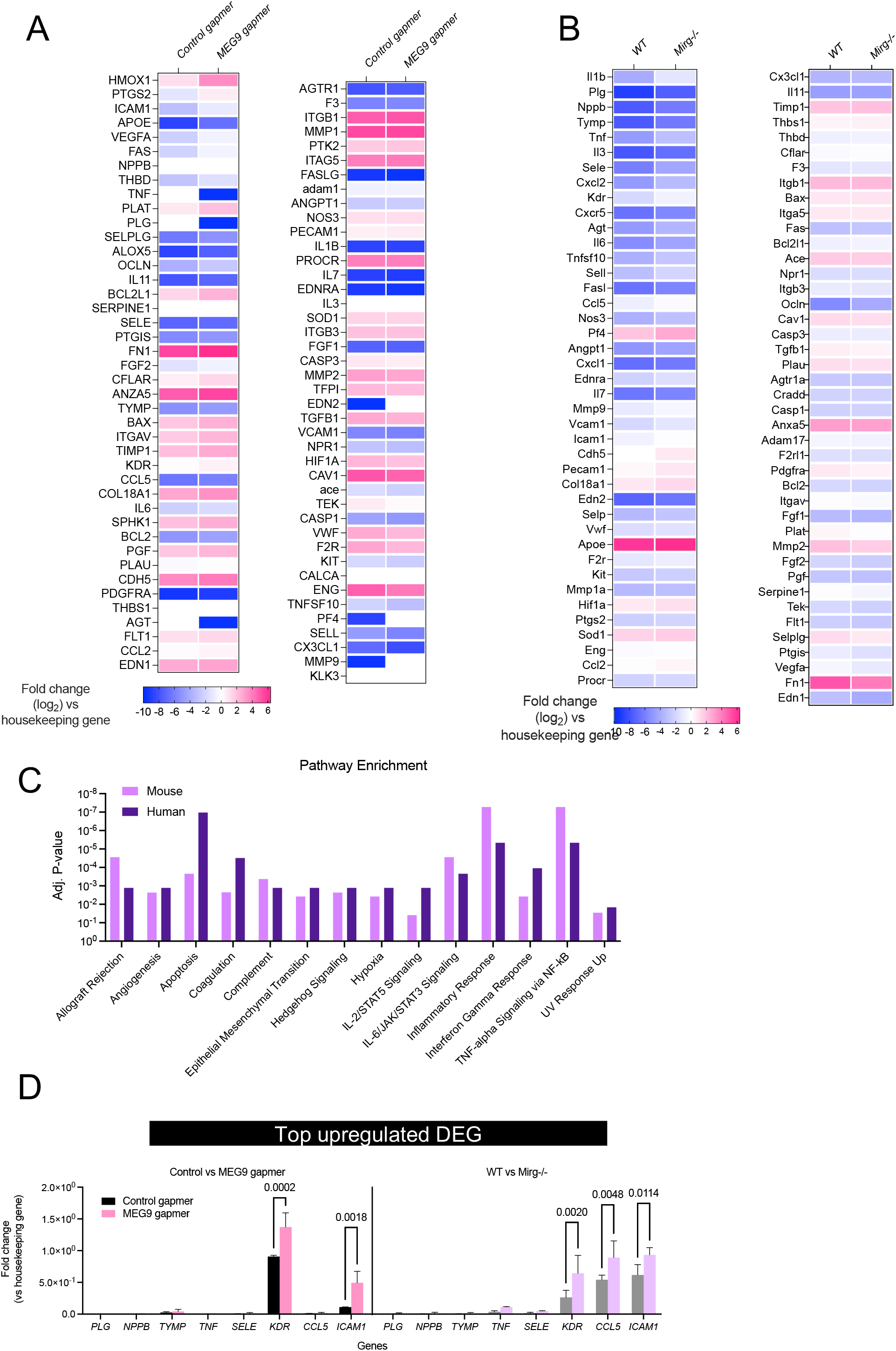
Identification of putative pathways regulated by MEG9/Mirg in vasculature. Heatmap depicting gene expression changes in A) HUVECs transfected with either a control GapmeR or MEG9 GapmeR B) Matrigel plugs from WT or Mirg-/-mice (n=2) from an angiogenesis qRT-PCR array. C) Pathway enrichment analysis from ENRICHR D) Top 8 differentially upregulated genes from the array. P-values from ANOVA with post-hoc Sidak test.

## Discussion

It is now widely appreciated that the majority of the human genome codes for non-protein-coding transcripts including lncRNAs and microRNAs. lncRNAs are thought to impact function by a variety of mechanisms including acting as a decoy (eg. GAS5), scaffold (HOTAIR), regulating splicing (MALAT1) etc [35]. Some specific lncRNAs have been identified as regulators of endothelial biology and angiogenesis [38, 39]. For instance, Tee et al showed that MALAT1 promotes tumor angiogenesis in response to hypoxia in part by inducing the secretion of FGF-2 [40]. Another lncRNA that is epigenetically regulated named MANTIS, appears to be a positive regulator of angiogenesis that interacts with the chromatin remodeling protein Brahma-like gene 1 (BRG1) [41]. Similarly, an EC enriched lncRNA STEEL was recently shown to be important for the formation of blood vessels by upregulating the transcription of eNOS and KLF2 [42]. Interestingly, some lncRNAs such as PUNISHER appear to be secreted in endothelial microvesicles and modulate angiogenesis distally highlighting a potentially non-cell autonomous mechanism of lncRNA function in cardiovascular pathology [43]. In fact, these lncRNAs together with a group of lncRNAs that regulate angiogenesis have been termed Angio-LncRs [35]. Using a screen for known lncRNAs, we sought out lncRNAs that were differentially regulated among stressors and stimuli and discovered MEG9 was a stress induced and growth factor suppressed lncRNA (Fig 1). Our work here adds MEG9 to the short list of lncRNAs that regulate angiogenesis in vitro and in vivo.

MEG9 resides on a unique imprinted locus on the genome – the Dlk1-Dio3 cluster [29]. The Dlk1-Dio3 cluster on Chr 14 in humans and Chr 12 in mice, harbors more than 50 microRNAs, several lncRNAs, and an array of snoRNAs. Several of these transcripts appear to have specific roles in cardiovascular development and angiogenesis [31]. For example, miR-300 is a negative regulator of cardiomyocyte progenitor differentiation [44]. Other miRs such as miR-410, miR-495 have been implicated in cardiac hypertrophy and high-fat diet induced cardiac remodeling [45, 46]. Among the lncRNAs, Gtl2/MEG3 appears to be an abundantly expressed lncRNA in endothelial cells and has complex roles in endothelial function, including regulation of DNA damage response and cellular senescence [25, 47]. While the cluster organization in the locus is conserved in all mammals, there are some differences between the mouse and human loci. The mouse Mirg locus harbors miRs, whereas the human MEG9 does not harbor any miRs. It has been shown that the ncRNAs in this locus are expressed from the non-methylated maternal allele in contrast to the protein coding genes that are predominantly expressed from the paternal allele.

We show that while MEG9 functions as a suppressor of EC death in vitro (Fig 2), the in vivo phenotype appears to be complex and sex-specific (Fig 3). Given the expression of the cluster lncRNAs in females, it is not surprising that we observed more vascular leak and less platelets in the female mice. However, male mice also showed a slight increase in lymphocytes that remains to be characterized (Fig 3). Our observations complement the findings from a comprehensive work on the non-coding RNA landscape of human hematopoiesis that found that lncRNAs in the Dlk1-Dio3 cluster specifically regulate megakaryopoiesis. In fact, knockdown of three specific lncRNAs MEG3, MEG8, MEG9 all interfered with megakaryopoiesis in culture highlighting our findings of less platelets in female mice might be directly due to the mouse orthologs of these lncRNAs.

Kamesmaran et al confirmed how susceptible the DLK1-DIO3 cluster is to methylation by targeting the specific promoter of lncRNA MEG3 in islets from type 2 Diabetes patients [48]. We have observed in our profiles that MEG9 and MEG3 respond in opposite directions to DNA damage and growth factor responses. While MEG3’s role in vascular pathology has been broadly described, the role and function of MEG9 is still unknown. Interestingly, Boon et al have described MEG3 was significantly upregulated in senescent HUVECs and in human cardiac atria from aged patients’ samples. Loss of function studies with MEG3-LNA increased sprouting angiogenesis *in vitro* and perfusion *in vivo* [49] and [50]. In myocardial infarction, MEG3 was also upregulated and a trigger for cell death and cardiomyocyte apoptosis [51].

These studies together with our results point out two lncRNAs encoded in the same megacluster are epigenetically regulated through DNA methylases and with opposite and complementary modulators. Whether or not MEG9 and MEG3 balance each other inducing and regulating opposite pathways in angiogenesis needs further investigation. At this point, we can conclude is MEG9 inhibition *in vitro* leads to endothelial cell death and decreases sprouting angiogenesis whereas the broader loss of the Mirg cluster results in sex-specific vascular leak and decrease in platelets.

A brief investigation of common pathways between the mouse and human loss of function genotypes yielded the VEGF receptor KDR, and cell adhesion molecule ICAM-1 as two genes upregulated in both species (Fig 4). ICAM-1 has been shown to impact platelet activation and increase vascular permeability in response to radiation [52]. Given some of the previous studies of VEGR-2 inhibitors in combination with genotoxic agents, Cisplatin and Gemcitabine show aberrant clotting events [53], our findings of KDR upregulation with the MEG9 loss of function are intriguing. Our data also highlighted Endothelin-1 as a protein that was significantly downregulated with a MEG9 knockdown (Supplementary Fig 5). Endothelin-1 is known to be a potent vasoconstrictor with diverse roles in EC pathophysiology. Interestingly, early studies showed an impact of Endothelin-1 on vascular permeability and cross-talk with the platelet-activating factor pathways [54]. While our mechanistic studies are limited in scope, we propose that further investigation into MEG9, regulation of its expression, and its function in vascular cells will elucidate the biological mechanism(s) by which this lncRNA affects endothelial cells.

## Acknowledgements

This work was supported by funding from NHLBI to S.A. (R01 HL137779 and R01 HL143803). We thank Marlee Feltham, Rishima Mukherjee, and Mallorie Mitchem for technical help and members of the Anand lab for valuable discussions. C.E-D. is supported by a training grant from NHLBI 5T32HL129964-07. A.B is supported by a training grant from NIGMS 5T32GM142619-02 and funds from the Knight Cancer Institute. The Genotype-Tissue Expression (GTEx) Project was supported by the Common Fund of the Office of the Director of the National Institutes of Health, and by NCI, NHGRI, NHLBI, NIDA, NIMH, and NINDS. The data used for the analyses described in this manuscript were obtained from the GTEx Portal Release V8 on 12/05/22 and dbGaP Accession phs000424.v8.p2

## Author Contributions

C. E-D. and S.A designed experiments, E. F-B, S.K., R.W., A.B., R.R., C.E-D performed experiments, analyzed the data, C.E-D. and S.A wrote the manuscript.

