## Supplementary Figures 1-7 for "DNA damage-induced lncRNA MEG9 impacts angiogenesis"

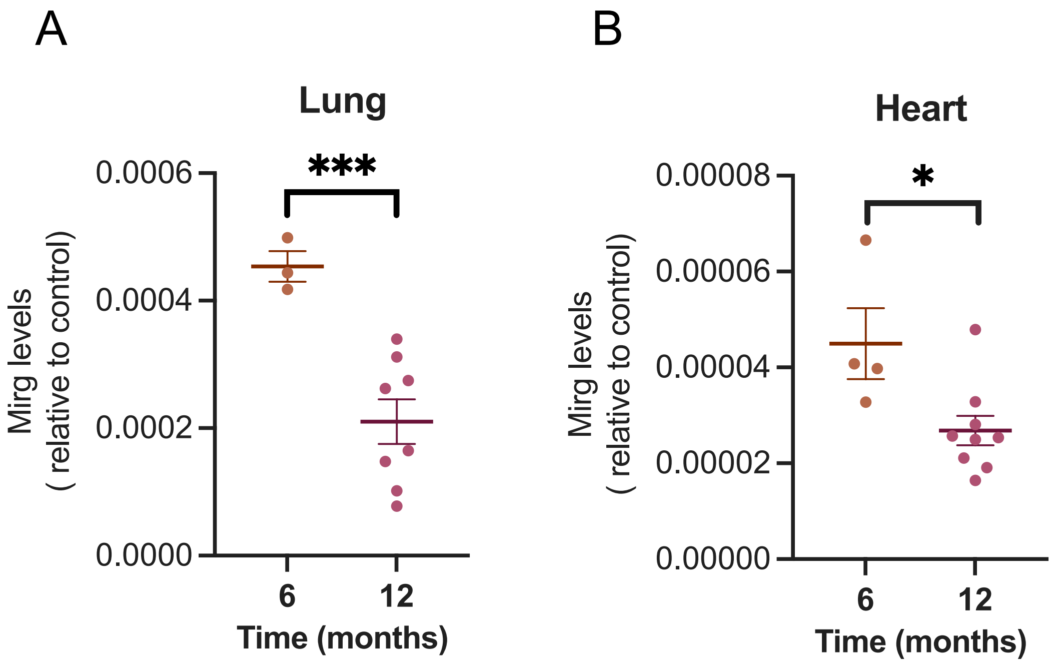


**Supplementary Figure 1. Mirg expression in mouse tissues.** (B) or Hearts (C) of 6 month or 12 old mice (equal numbers of male and female mice) Mirg levels were measured using qRT-PCR. *P<0.05 by Student's T-test.

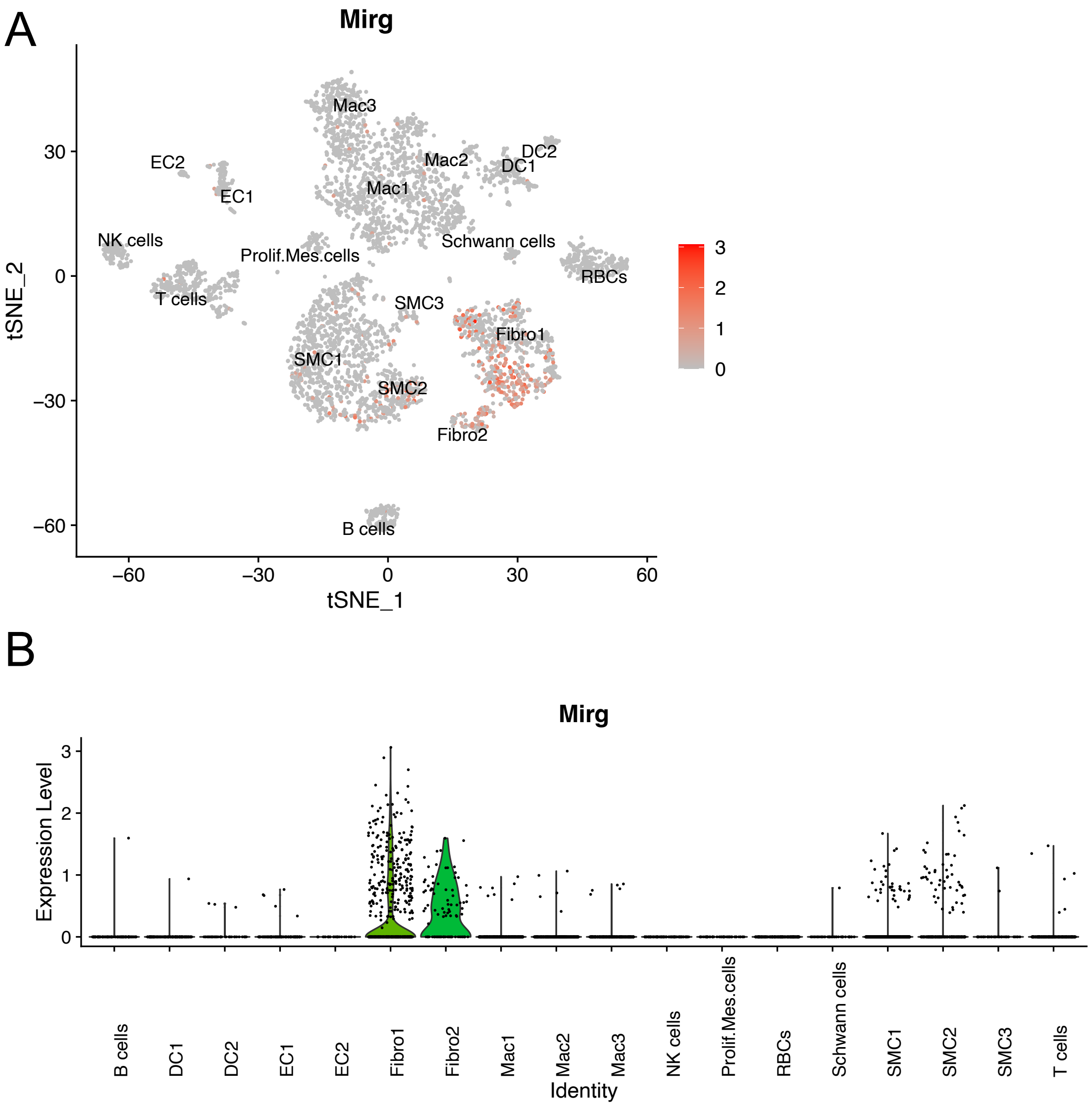


**Supplementary Figure 2. MEG9 expression in different cell populations in the aorta.** (A) tSNE plot and (B) expression for Mirg in mouse aorta from scRNAseq datasets ([GSE152583]) analyzed from the Cardiovascular Atlas (CLARA) portal.

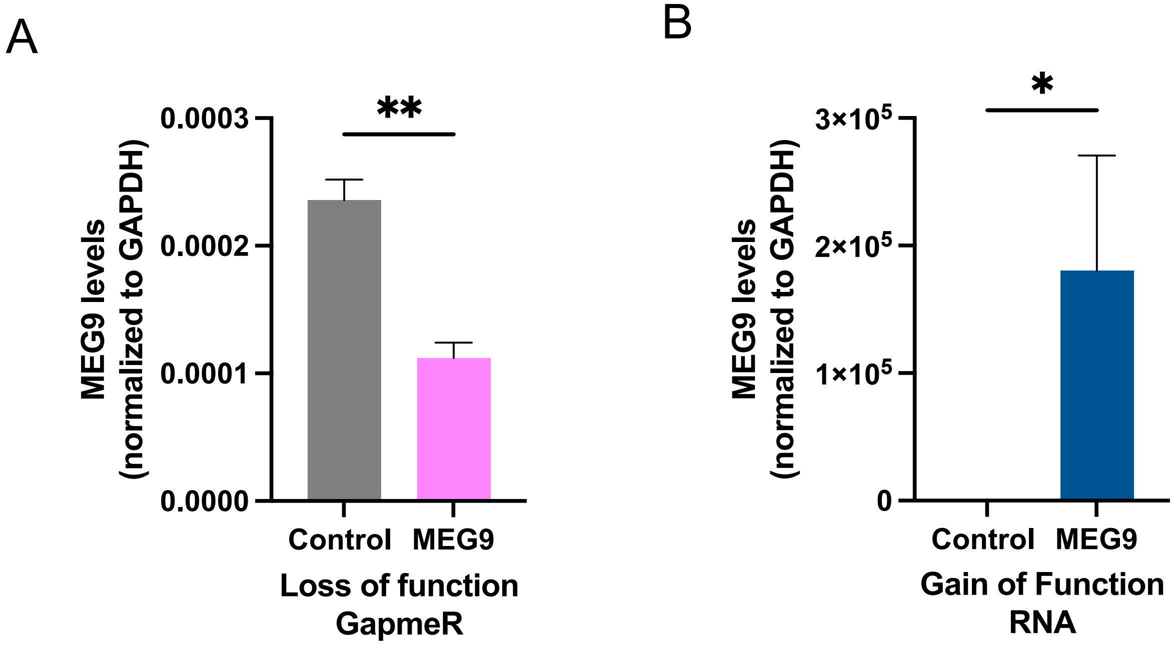


**Supplementary Figure 3. Validation of tools to modulate MEG9 expression.** A) HUVECs were transfected with siRNA Gapmer against MEG9 or control Gapmer B) in vitro transcribed MEG9 RNA or a control RNA. MEG9 mRNA levels were measured by qRT-PCR 24h post-transfection. *P<0.05, ** P<0.01 from a two-tailed Student’s T-test.

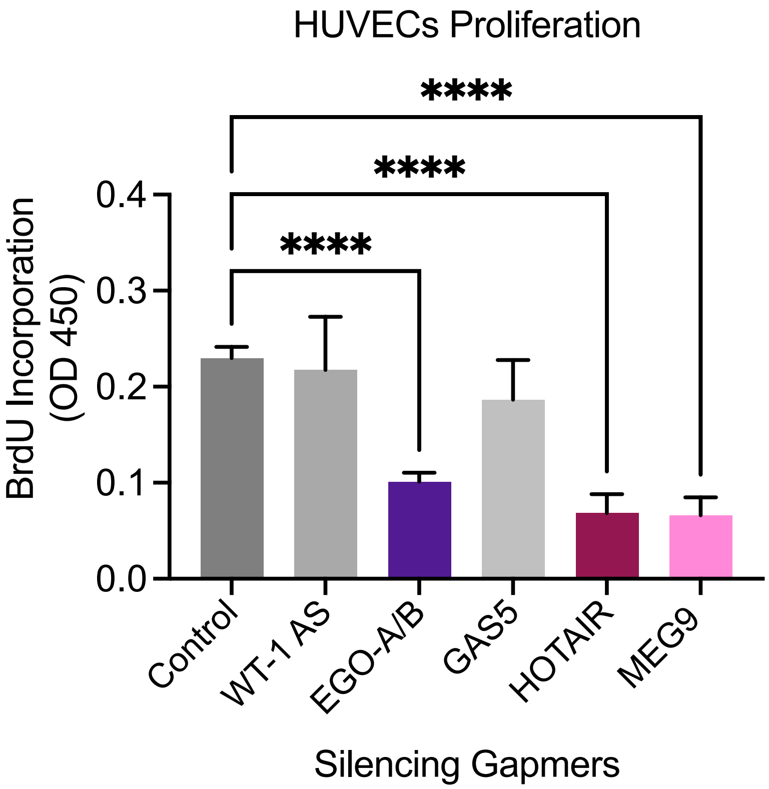


**Supplementary Figure 4. Function of top-ranked lncRNAs in HUVECs.** A) HUVECs were transfected with siRNAs Gapmer against indicated target lncRNAs. Cells were pulsed with BrdU 30h post transfection and BrdU incorporation was measured by ELISA 48h post transfection. One of two independent experiments. **** indicates P<0.0001 by ANOVA with post-hoc Dunnett's test.

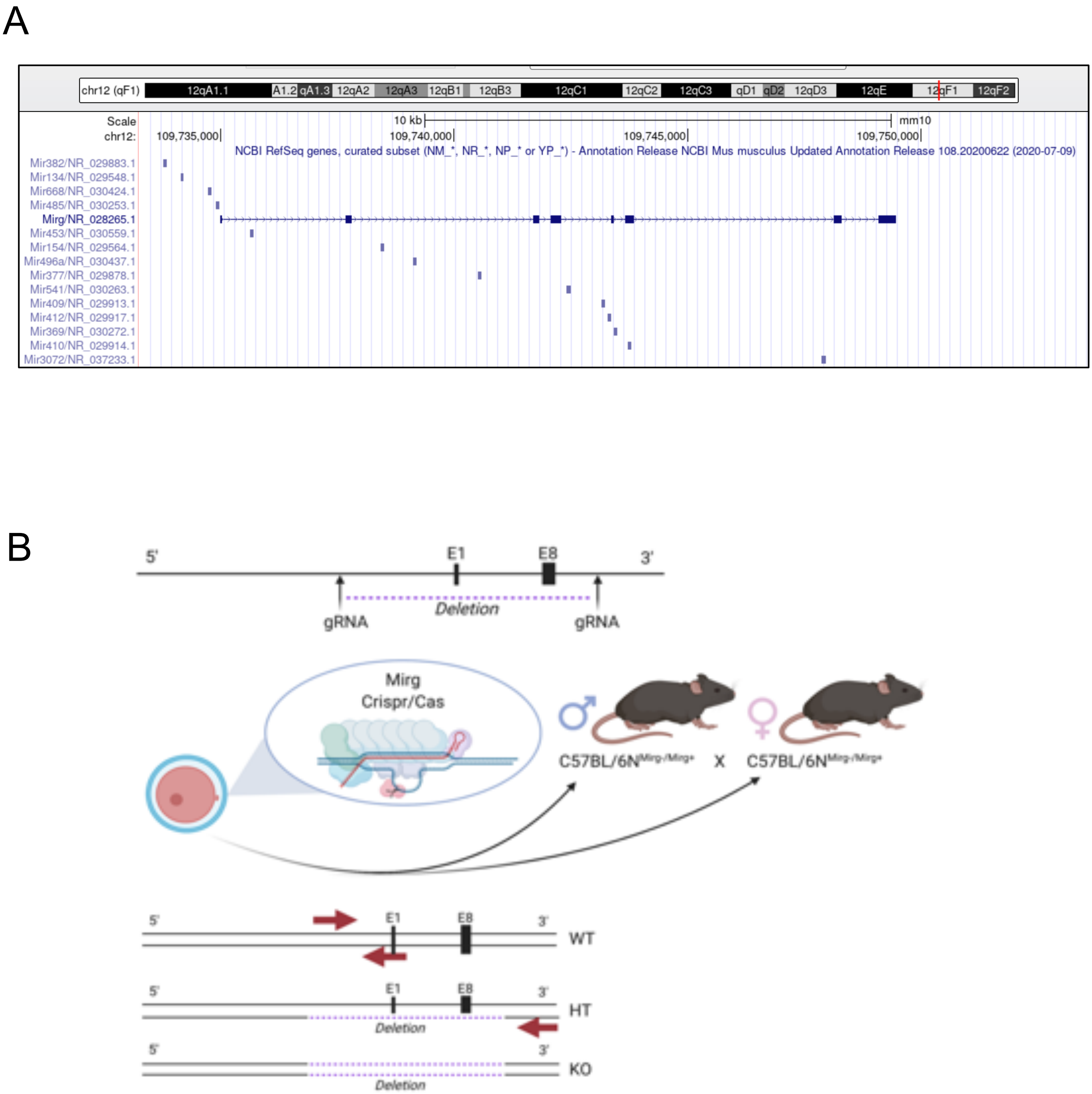


**Supplementary Figure 5. Schematic depicting CRISPR strategy for deletion of Mirg locus in mice.**

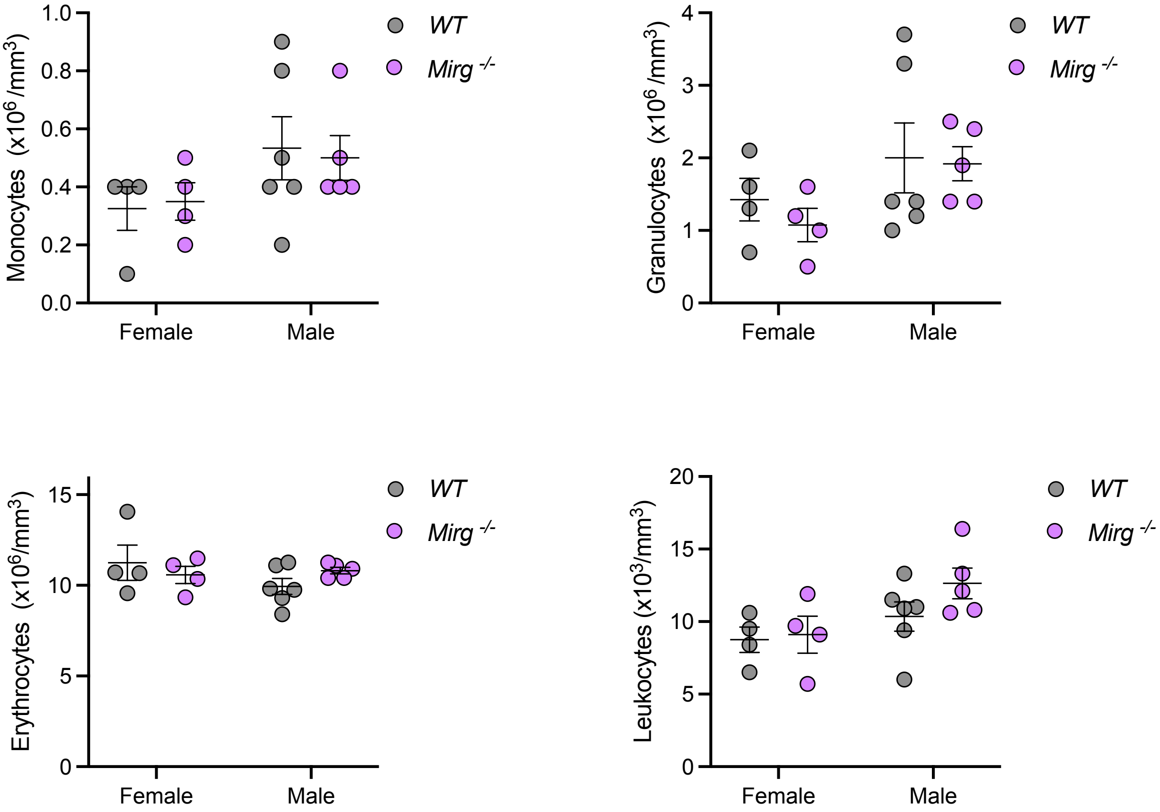


**Supplementary Figure 6. Blood counts of Mirg-/- mice do not show alterations in several major cell types.** Counts of indicated cell types from WT vs Mirg-/- mice.

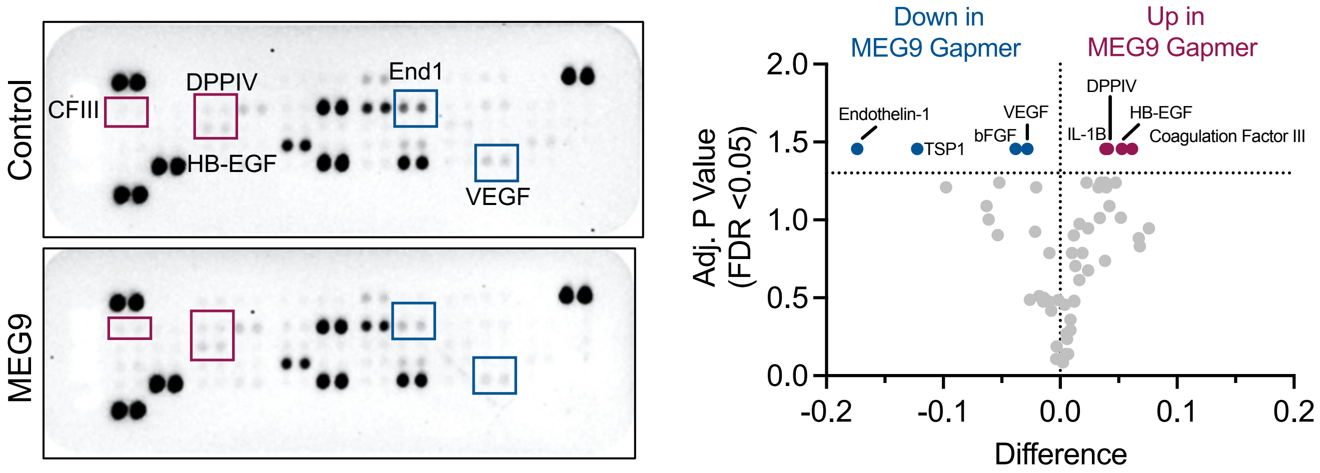


**Supplementary Figure 7. Knockdown of MEG9 impacts angiogenic signaling pathway proteins.** HUVECs were transfected with siRNA Gapmer against MEG9 or control Gapmer. 48h later, lysates were run on a Proteome Profiler Western blot membrane array with 55 different capture antibodies. Images were quantified using Image J, normalized to the reference spots. Volcano plot shows differences in the MEG9 pixel intensities vs the adjusted P-values calculated by false discovery rates.
